# Functional analyses of two novel *LRRK2* pathogenic variants in familial Parkinson’s disease

**DOI:** 10.1101/2021.12.03.470891

**Authors:** I Coku, E Mutez, S Eddarkaoui, S Carrier, A Marchand, C Deldycke, L Goveas, G Baille, M Tir, R Magnez, X Thuru, G Vermeersch, W Vandenberghe, L Buée, L Defebvre, B Sablonnière, MC Chartier-Harlin, JM Taymans, V Huin

## Abstract

**Background:** Pathogenic variants in the *LRRK2* gene are a common monogenic cause of Parkinson’s disease. However, only seven variants have been confirmed to be pathogenic.

**Objectives:** We identified two novel *LRRK2* variants (H230R and A1440P) and performed functional testing.

**Methods:** We transiently expressed wildtype, the two new variants, or two known pathogenic mutants (G2019S and R1441G), in HEK-293T cells, with or without LRRK2 kinase inhibitor treatment. We characterized the phosphorylation and kinase activity of the mutants by western blotting. Thermal shift assays were performed to determine the folding and stability of the LRRK2 proteins.

**Results:** The two variants were found in two large families and segregate with the disease. They display altered LRRK2 phosphorylation and kinase activity.

**Conclusions:** We identified two novel *LRRK2* variants which segregate with the disease. The results of functional testing lead us to propose these two variants as novel causative mutations for familial Parkinson’s disease.

## INTRODUCTION

Parkinson’s disease (PD) is a neurodegenerative disorder characterized by the selective loss of dopaminergic neurons from the *substantia nigra pars compacta* associated with Lewy bodies rich in aggregated alpha-synuclein and lipids in surviving neurons (1). Most cases are sporadic. However PD can be concentrated in certain families and/or have an early-onset (≤ 45 years). It can be caused by a monogenic form of the disease explaining < 10% of familial cases and a still lower frequency of apparently sporadic cases (2–4).

Pathogenic variants in the leucine-rich repeat kinase 2 (*LRRK2*) gene are among the most common genetic causes of familial and sporadic PD. Indeed, the G2019S pathogenic variant is the most frequent, with its prevalence reaching up to 29% in Ashkenazi Jewish and 37% in North African Berber populations (5). More than 80 rare coding sequence variants in *LRRK2* have been reported to be linked to PD thus far, but only seven (*i*.*e*., N1437H, R1441G, R1441C, R1441H, Y1699C, G2019S, and I2020T) (Fig. 1A) have been confirmed to be pathogenic and responsible for PD with a Mendelian inheritance (5,6). To date, all pathogenic variants are located in the kinase or Roc-COR domains.

**Figure 1.**
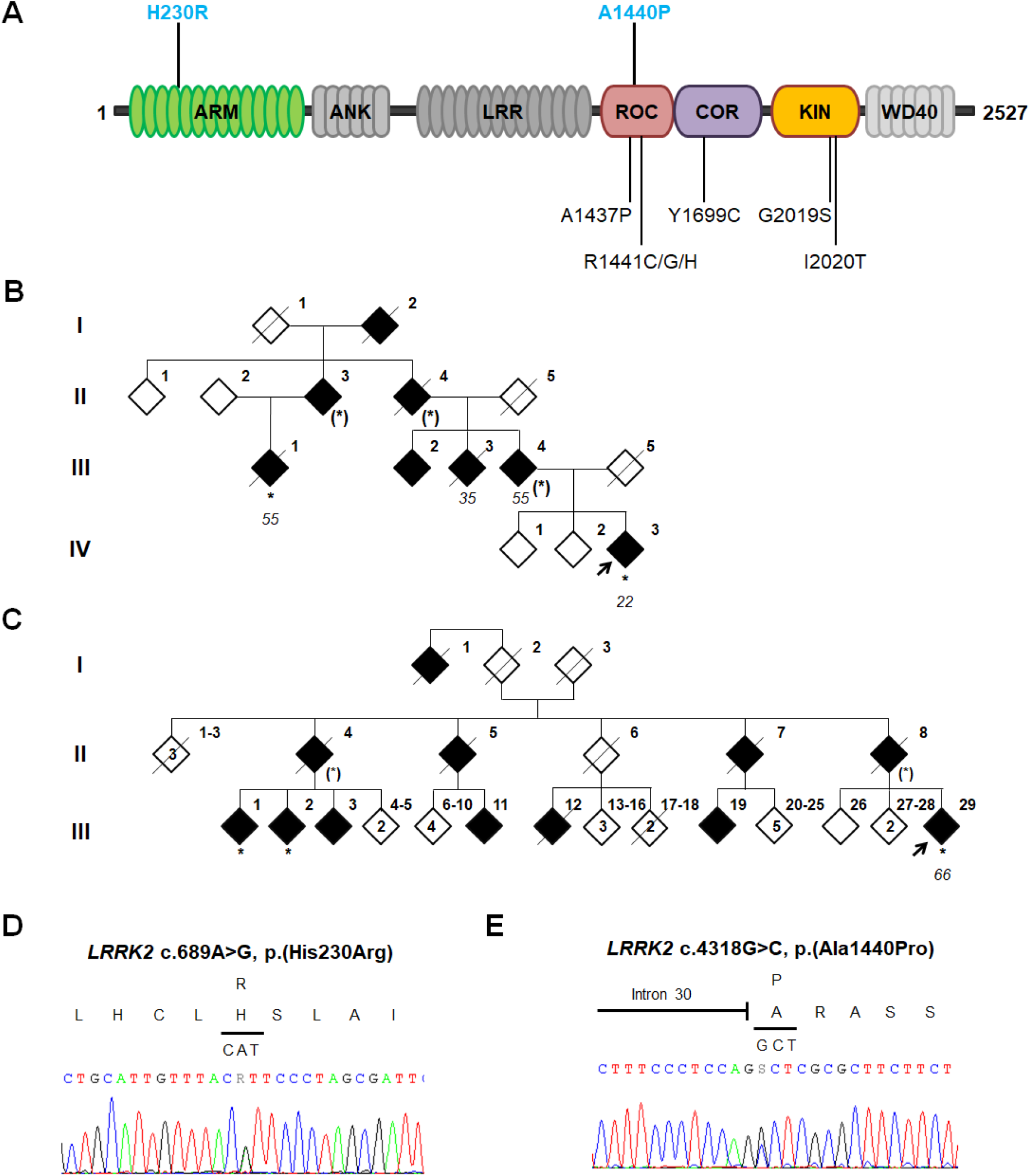
Nature and position of two novel *LRRK2* pathogenic variants. **A**. Schematic linear representation of LRRK2 protein. The two novel pathogenic variants are indicated in bold above the protein and the seven known pathogenic mutants are indicated below the protein. Each domain of LRRK2 is named: ARM, armadillo repeats; ANK, ankyrin repeats; LRR, leucine-rich repeats; ROC, Ras of complex proteins GTPase; COR, C-terminal of ROC; KIN, kinase; WD40, WD40 repeats. **B**. Family tree of family 1, with the *LRRK2* H230R variant. **C**. Family tree of family 2, with the *LRRK2* A1440P variant. The probands are denoted by a black arrow. Filled black symbols denote clinically affected members and open symbols indicate unaffected individuals. / = deceased, * = genotyped carrier, (*) = obligatory carrier. The numbers under each individual correspond to the age of onset of PD. Family pedigrees have been anonymized for confidentiality. **D-E**. Electropherograms of heterozygous pathogenic variants of *LRRK2* for NM_198578.3:c.689A>G, p.(His230Arg) **(D)** and NM_198578.3:c.4318G>C, p.(Ala1440Pro) **(E)**.

The major hypothesis to explain the pathophysiology of *LRRK2* pathogenic variants in PD is a gain of function that induces an increase in kinase activity and hyperphosphorylation of the substrate proteins (7). Altered autophosphorylation of serine 1292 and increased Rab proteins phosphorylation have been observed in *LRRK2* pathogenic variants *in cellulo* and are indicators of kinase activity (8).

Here, we identified two novel LRRK2 variants: H230R in the armadillo domain and A1440P in the ROC domain. We tested the kinase activity of these new variants *in cellulo* and assessed their thermal stability to demonstrate their pathogenicity.

## MATERIALS AND METHODS

### Subjects

Two families displaying PD with an autosomal-dominant inheritance pattern were screened during targeted next-generation sequencing of PD genes in a diagnostic setting (Fig. 1B-C). Patients underwent a detailed clinical evaluation in the department of Neurology and Expert Center for Parkinson’s disease at the Lille, Amiens, Bruges or Leuven Hospitals. Clinical diagnoses were reviewed according to the international diagnostic criteria for PD (9). Extensive genetic analyses were performed to eliminate other genetic diseases (Supplementary material). All individuals gave their written informed consent. The study was conducted according to the French ethics regulations (Lille ethics committee, Protocole Convergence, CPP/2008/009).

### Functional testing

Briefly, we used a previously described plasmid construct (10) for the wildtype (WT) human LRRK2 (pLV-CSJ-3FLAG-LRRK2-WT) as a template to introduce the two novel variants (H230R and A1440P) and two known pathogenic variants (G2019S and R1441G) as positive controls. We transiently expressed WT, or the LRRK2 mutants, in HEK-293T (human embryonic kidney cells that express the SV40 large T antigen) cells, with or without with LRRK2 kinase inhibitor (11). We performed western blot to asses LRRK2 hetero- and auto-phosphorylation, and to characterize the phosphorylation of a known LRRK2 substrate, RAB10 at threonine 73. Lastly, we purified the LRRK2 proteins and performed thermal shift assay.

Additional methods are described in the Supplementary material.

## RESULTS

### Genetic analyses

Next-generation sequencing revealed two missense variants in *LRRK2* (NM_198578.3): c.689A>G, p.(His230Arg) in exon 6 in family 1 (Fig. 1B,D), and c.4318G>C, p.(Ala1440Pro) in exon 31 in family 2 (Fig. 1C,E). There were no pathogenic variants in the other PD genes. These variants were absent from databases of healthy individuals (gnomAD v3.1.1) (12). The variant A1440P is located in a mutational hotspot in the Roc domain and multiple prediction tools (DANN, MutationTaster, FATHMM, GERP++, LRT, MetaLR, MetaSVM, and PROVEAN) favored a deleterious effect. Segregation analyses showed the variant A1440P to be heterozygous in two affected cousins of the proband and the variant H230R to be heterozygous in the proband’s second cousin affected with PD. Co-segregation analyses in these two families provided a moderate level of evidence of pathogenicity (13). Details of clinical phenotypes are provided in Supplementary data.

### Kinase activity

We first studied the phosphorylation of LRRK2 by other kinases at serine 910 and 935 (Fig. 2A-D). In HEK-293T cells transiently expressing WT or mutant forms of LRRK2, WB analyses showed a higher phosphorylation rate for G2019S than for WT at serine 910 (1.6-fold increase, *p* = 0.005) and serine 935 (two-fold increase, *p* = 0.016); whereas the R1441G mutant showed two-fold lower phosphorylation at serine 910 (*p* = 0.009) and no difference from the WT at serine 935 (*p* = 0.215), as expected (14). The H230R mutant showed a slight but nonsignificant increase in phosphorylation at serine 910 (*p* = 0.561) and a 2.4-fold increase at serine 935 (*p* = 0.003), whereas the mutant A1440P showed decreased phosphorylation at serine 910 (*p* = 0.003) and a slight but nonsignificant decrease in phosphorylation at serine 935 (*p* = 0.319).

**Figure 2.**
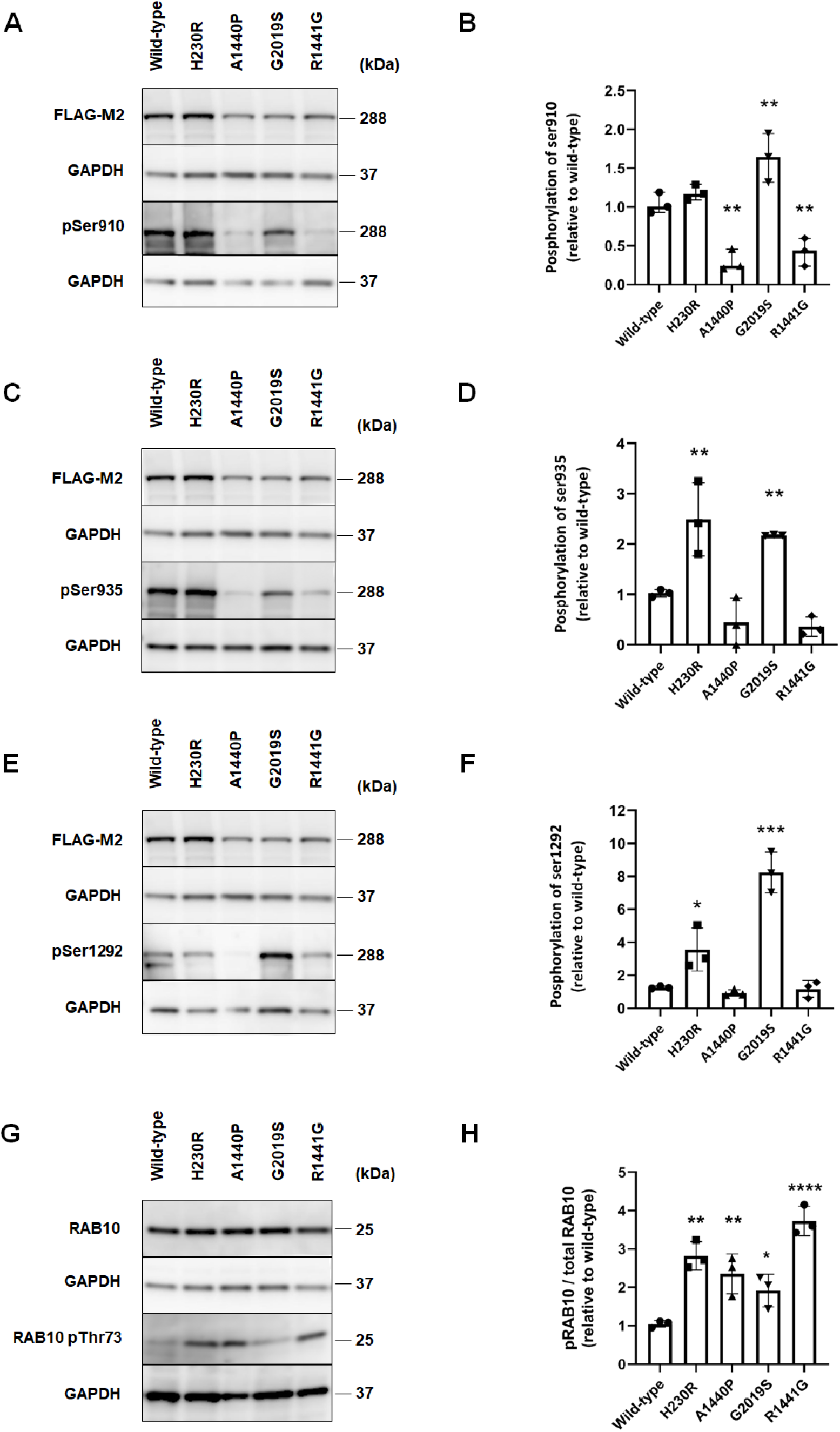
Comparison of Ser910/Ser935/Ser1292 phosphorylation sites in PD-associated mutants. Representative western blots and quantification of LRRK2 phosphorylation at serines 910 **(A-B)**, 935 **(C-D)**, and 1292 **(E-F)** in WT and LRRK2 mutants. Representative western blot **(G)** and quantification **(H)** of RAB10 phosphorylation at threonine 73 for the WT and LRRK2 mutants. Error bars indicate the standard deviation of replicates (n = 3). kDa = Kilodalton. * *p* < 0.05, ** *p* < 0.01, *** *p* < 0.001; **** *p* < 00001.

We then studied the phosphorylation of serine 1292 (Fig. 2E-F), which is an indicator of LRRK2 autophosphorylation and kinase activity. As expected, we observed higher phosphorylation of serine 1292 for the mutant G2019S (≈ eight-fold increase, *p* < 0.0001), whereas R1441G showed no difference relative to WT (*p* = 0.997) (15). The mutant H230R showed 3.5-fold greater phosphorylation at serine 1292 (*p* = 0.013), whereas A1440P showed no difference relative to WT (*p* = 0.999).

We next characterized the phosphorylation of a known LRRK2 substrate, RAB10 at threonine 73. We observed two-fold higher phosphorylation for G2019S (*p* = 0.046) and 3.7-fold higher phosphorylation for R1441G (*p* < 0.0001). The rate of phosphorylation of RAB10 was approximately 2.8-fold higher for H230R and 2.3-fold higher for A1440P (*p* < 0.001 and *p* = 0.005, respectively) (Fig. 2G-H).

### LRRK2 inhibitor

Finally, we investigated the kinase activity of our mutants using MLi-2, a highly selective LRRK2 kinase inhibitor and observed how the inhibition of LRRK2 protein kinase affects the phosphorylation of LRRK2 or its substrate RAB10. We compared HEK-293T cells transiently expressing WT or mutant forms of LRRK2 and treated with 100 nM MLi-2 or 0.01% DMSO for 1 h (Supplementary Fig. S1). Under all conditions, the cells treated with MLi-2 showed less mean phosphorylation than those treated with DMSO. There was significantly less phosphorylation of serine 910 for G2019S (*p* = 0.013), serine 935 for H230R (*p* = 0.003) and G2019S (*p* = 0.032), and serine 1292 for H230R (*p* = 0.042) and G2019S (*p* = 0.006). Treatment with MLi-2 almost completely suppressed the phosphorylation of RAB10 at threonine 73 under all conditions (Supplementary Fig. S2).

### Thermal stability

The H230R and A1440P variant proteins did not show notable differences in their respective profiles, Ti or Δ Ratio relative to the WT protein. The initial ratio, which indicates the level of folding at basal condition, was similar for H230R (0.34 ± 0.019; *p* = 0.97) and lower for A1440P (0.31 ± 0.011; *p* = 0.036), suggesting that these proteins are correctly folded and stable (Supplementary data; supplementary Fig. S3).

## DISCUSSION

We identified two novel pathogenic variants of the *LRRK2* gene from large autosomal-dominant PD families. Our segregation analyses and the characteristics of the variants (frequency in databases, location in the gene, prediction tools) provide sufficient evidence to consider them to be at least “likely pathogenic” for A1440P according to the ACMG guidelines for the classification of genetic variants (16). Subsequent functional analyses allowed considering these two variants as “pathogenic”.

*LRRK2* A1440P shows phosphorylation rates at serine 910, 935, and 1292 similar to those of R1441G located at the adjacent position (14,15). On the other hand, H230R shows a pattern of phosphorylation more similar to that of G2019S (14,15). All previously reported pathogenic LRRK2 mutants show greater phosphorylation of Rab proteins (7,17). Similarly, we observed approximately two-fold greater phosphorylation of RAB10 at threonine 73 for both mutants than for WT LRRK2. These results are comparable with those of previous studies, suggesting that LRRK2 kinase activity cannot be uniformly predicted by its autophosphorylation and cellular phosphorylation site status (18).

Given its location and the similar phosphorylation pattern of the mutated protein, the pathological effects of A1440P are likely to be similar to those of other pathogenic variants located in the Roc-COR domain, such as R1441G. However, the pathophysiology and increased kinase activity of the H230R variant, located in the ARM domain, is less obvious. Another reported variant in this domain, A211V, also showed a slight increase in kinase activity (19,20). Kishore et al. identified A397T, G472R, and L550W mutations but they did not describe the kinase activity of these rare variants (21). Another rare variant, N551K, belonging to a protective haplotype (N551K-R1398H-K1423K) has been reported for PD patients (22,23) but the mechanisms explaining how this haplotype confers neuroprotection in PD is not clear and it has not been functionally assessed. Only one transcriptomic study in Drosophila melanogaster has identified altered pathways associated with N551K, including alterations of the oxidoreductase pathway (24). Structural analysis of full-length human LRRK2 has shown that the ANK and LRR domains interact with the kinase domain but not the ARM domain, which shows flexibility relative to the rest of the protein (25). Rab proteins directly interact with LRRK2 via the ARM domain (26,27) but the H230R variant is not located in the potential Rab-interacting regions of this domain (residues 386–392) (25). It has also been suggested that amino-acid substitutions of the conserved ARM domain of LRRK2 enhance interactions with FADD and induce apoptosis via caspase 8 (28). Another study reported that LRRK2 interacts with Hsp90 via its ARM domain and then Hsp90 subsequently interacts with the E3 ubiquitin ligase CHIP to decrease LRRK2 CHIP-mediated degradation (29). The ARM domain interacts with RAB7L1 (RAB29) (27), a membrane-anchored RAB GTPase that recruits LRRK2 to the trans-Golgi network or lysosomes via the ANK domain and highly stimulates its kinase activity (27).

In conclusion, we have identified two novel *LRRK2* variants, H230R and A1440P, which segregate with the disease in large PD families. We show that H230R and A1440P alter the phosphorylation rates of LRRK2 and its ability to phosphorylate its substrate RAB10. Further studies on these rare pathogenic variants should help us to better understand how LRRK2 dysfunction causes PD and may have implications for future treatment strategies against *LRRK2*-related disorders.

## ACKNOWLEDGEMENTS

We are grateful to the patients and family members for their participation in this study.

## DATA AVAILABILITY STATEMENT

The data that support the findings of this study are available from the corresponding authors upon reasonable request.

## AUTHORS’ ROLES

I.C. = Conception, Execution, Data curation; Statistical Analysis: Execution; Manuscript: Writing of the first draft.

E.M. = Organization, Execution, Data curation; Manuscript: Review, and Critique.

S.E. = Execution, Data curation

S.C. = Execution, Data curation

A.M. = Execution, Data curation

C.D. = Execution, Data curation

L.G. = Execution, Data curation

G.B. = Execution, Data curation

M.T. = Execution; Manuscript: Review and Critique.

R.M. = Execution, Data curation; Manuscript: Writing of the first draft.

X.T. = Organization; Manuscript: Review and Critique.

G.V. = Data curation; Manuscript: Review, and Critique.

W.V. = Data curation; Manuscript: Review, and Critique.

L.B. = Organization; Manuscript: Review and Critique.

L.D. = Organization; Manuscript: Review and Critique.

B.S. = Organization, Manuscript: Review and Critique.

M-C.C-H. = Conception, Organization; Statistical Analysis: Review and Critique; Manuscript: Review and Critique.

J-M.T. = Conception; Statistical Analysis: Design, Review, and Critique; Manuscript: Review and Critique.

V.H. = Conception, Data curation; Organization; Statistical Analysis: Design, Review, and Critique; Manuscript: Writing of the first draft, Review, and Critique.

