## Supplemental Files for "Functional analyses of two novel *LRRK2* pathogenic variants in familial Parkinson’s disease"

### SUPPLEMENTARY MATERIAL

#### Genetic screening

Genomic DNA was isolated from peripheral blood according to standard procedures. Multiplex ligation-dependent probe amplification (MLPA) and targeted next-generation sequencing of 72 Parkinsonism-related genes were performed in the probands.

*Multiplex ligation-dependent probe amplification (MLPA)*

Genomic DNA was isolated from peripheral blood according to standard procedures. We performed MLPA for all patients using the P051 and P052 SALSA MLPA probe mixes for the molecular diagnosis of inherited Parkinson's disease (MRC Holland, Amsterdam, The Netherlands) according to the manufacturer's instructions. Electrophoresis of the MLPA products was performed on an ABI 3730 DNA analyzer (Applied Biosystems, Foster City, CA, USA). We analyzed the MLPA data using GeneMapper 2.5 (Applied Biosystems, Foster City, CA, USA) and COFFALYSER (MRC Holland, Amsterdam, The Netherlands) software.

*Targeted sequencing*

Exon capture of the 72 selected genes and libraries was prepared using the Haloplex kit (Agilent Technologies, Santa Clara, CA, USA) according to the manufacturer's instructions. Pooled libraries (n = 8) prepared using the Parkinson's disease genes panel were sequenced using paired-end, 150-cycle chemistry on an Illumina MiSeq (Illumina, San Diego, CA, USA) using MiSeq Reageant kit v2 according to the manufacturer's protocol. All variants were confirmed by Sanger sequencing. All genetic variants were confirmed by Sanger sequencing.

*List of the 72 selected genes*

The following genes were analyzed by high throughput sequencing: *ADH1C, AFG3L2, AP5Z1, ATP13A2, ATP1A3, ATP6AP2, ATP7B, C10ORF2, C19ORF12, CHCHD10, CHCHD2, CLN3, COASY, COQ2, CP, CRAT, CSF1R, DCAF17, DCTN1, DNAJC12, DNAJC13, DNAJC5, DNAJC6, EIF4G1, FBXO7, FA2H, FTL, GBA, GCH1, GIGYF2, GRN, HTRA2, KCND3, KIF5A, LRRK2, MAPT, PANK2, PARK2, PARK7, PDE8B, PDGFB, PDGFRB, PINK1, PLA2G6, POLG, PRKRA, PRNP, PTS, RAB29, RAB39B, REPS1, SCP2, SLC18A2, SLC20A2, SLC30A10, SLC39A14, SLC6A3, SNCA, SPG11, SYNJ1, TAF1, TH, TMEM240, TTC19, UBTF, UCHL1, VPS13A, VPS13C, VPS35, WDR45, XPR1, ZFYVE26.*

*Bioinformatics analysis*

The sequence data were aligned to the hg19 assembly version of the human genome using BWA v0.7.17 (1). Variant calling, joint genotyping, and recalibration were performed using GATK v3.7 (2). Sequence analysis was performed using SeqPilot (v4.3.1, JSI Medical Systems, Ettenheim, Germany), MiSeqReporter (v2.6, Illumina, San Diego, CA, USA) and Genesearch (PhenoSystems, Braine le Chateau, Belgium) software. Variants were annotated and filtered using Variant Studio software (v3.0, Illumina, San Diego, CA, USA). Harmful effects were predicted using Alamut software (v2.9.0, Interactive BioSoftware, Rouen, France).

**_LRRK2_ expression constructs and site-directed-mutagenesis**

A previously described plasmid construct (Drouyer et al., 2021) of the wildtype (WT) human *LRRK2* cDNA with an N-terminal triple Flag epitope tag (pLV-CSJ-3FLAG-LRRK2-WT) was used as a template for all subsequent expression constructs. The use of the N-terminal triple Flag epitope tag is useful for protein purification and does not induce artefactual phenotypes in transfected cells (4). For the introduction of known and novel We introduce the two novels pathogenic PD-linked sequence variations, we used the QuikChange® II XL site-directed mutagenesis kit (Agilent Technologies, Santa Clara, CA, USA) according to the manufacturer's instructions. The primer sequences are given in Supplementary table S1. Plasmids subjected to mutagenesis underwent full sequencing (GATC Biotech, Zurich, Switzerland) to verify that the expected mutations were introduced and that the constructs were not otherwise altered.

**Cell culture, transfection, and treatment with LRRK2 kinase inhibitor**

HEK-293T were cultured in Dulbecco's Modified Essential Medium (Invitrogen, Carlsbad, CA, USA) supplemented with 10% fetal bovine serum (FBS), 1% Glutamax, and antibiotics (100 units/ml penicillin G and 100 µg/ml streptomycin) (Life Technologies, Carlsbad, CA, USA) at 37°C under 5% $CO_2$. Transfection of the cells with 3xFlag-LRRK2 plasmid DNA (wildtype and mutants) and the empty vector backbone, as a negative control, was performed using Lipofectamine 2000 Transfection Reagent (LIPO 2000) (Invitrogen, Carlsbad, CA, USA) with a 1:2 DNA:LIPO 2000 ratio. The transfection mixes were prepared by dissolving 30 µg DNA in 1.5 ml OPTI-MEM with 60 µl LIPO 2000. Mixtures were incubated at room temperature for 20 min and subsequently added to HEK-293T cells at 80% confluence in 15-cm diameter Petri dishes. Cells were washed once 6 h later with PBS. Cells were treated 48 h later with MLi-2 (MedChemExpress, Monmouth Junction, NJ, USA) (5), a highly selective LRRK2 kinase inhibitor (100 nM), or the solvent DMSO (0.01%) for 1 h or were left untreated (negative controls). As a negative control, we transfected HEK-293T cells with the empty vector backbone plasmid.

**Western-blotting**

HEK-293T cells were collected 48 h after transfection, washed once in PBS, and resuspended in cell lysis RIPA buffer (100 mM Tris base, 300 mM NaCl, 0.2% SDS, 1% sodium deoxycholate, 2% NP-40, protease inhibitor cocktail (Sigma-Aldrich, Saint-Louis, MI, USA). Samples were sonicated 20 times at 40 Hz and then incubated for 60 min at 4°C on a rotator. Protein concentrations were determined using the Pierce™ BCA Protein Assay Kit (Thermo Fisher Scientific, Waltham, MA, USA). Cell extracts were mixed with Tris-Glycine SDS sample buffer (Novex, Life Technologies, Carlsbad, CA, USA) and boiled at 100°C for 10 min. After loading 15 µg total protein and molecular-weight markers (Novex and Magic Marks, Life Technologies, Carlsbad, CA, USA) onto 4–20% Novex Tris-Glycine gels (Novex, Life Technologies, Carlsbad, CA, USA), electrophoresis was carried out by applying 165 V for 1.5 h in SDS running buffer (Tris-Glycine running buffer: 25 mM TRIS pH 8.6, 190 mM glycine, 1% SDS, Novex; Life Technologies, Carlsbad, CA, USA). Proteins were electrophoretically transferred onto a 0.45-µm nitrocellulose membrane (G&E Healthcare, Chicago, IL, USA) at 30 V for 90 min. The quality of the protein transfer was determined by reversible Ponceau Red coloration (0.2% Xylidine Ponceau Red and 3% trichloroacetic acid). Membranes were blocked in 25 mM Tris-HCl pH 8.0, 150 mM NaCl, and 0.1% Tween-20 (v/v) (TBS-T) supplemented with 5% (w/v) skimmed milk (TBS-M) or 5% (w/v) bovine serum albumin (TBS-BSA), depending on the antibody, for 60 min. Membranes were incubated with primary antibodies overnight at 4°C. After washing in TBS-T, membranes were incubated with secondary antibodies for 45 min at room temperature. Revelation was performed using ECL™Western-blot detection reagents (G&E Healthcare, Chicago, IL, USA). The antibodies used for western-blotting are listed in Supplementary Table S2.

**Protein purification**

LRRK2 protein was purified essentially as previously described (6,7). Briefly, HEK-293T cells expressing 3xFlag-LRRK2 were solubilized at 4°C in lysis buffer containing 20 mM Tris-HCl pH 7.5, 150 mM NaCl, 1 mM EDTA, 1% Triton X-100, and 10% glycerol. Lysates were incubated for 30 min at 4°C on a rotator and then centrifuged for 15 min at 15000 × g. Anti-Flag M2 agarose beads (F3165, Sigma-Aldrich, Saint-Louis, MI, USA) were equilibrated by washing three times in lysis buffer. Then, lysates containing 3xFlag-tagged protein were incubated with anti-Flag M2 agarose beads for 48 h at 4°C on a rotator. After five washes (3 with 25 mM Tris-HCl pH 7.5, 400 mM NaCl, 1%Triton; 2 with 20 mM Tris-HCl pH 7.5, 200 mM NaCl, 5 mM MgCl2, 1mM DTT, 0.02%Triton), proteins were eluted in elution buffer (20 mM Tris-HCl pH 7.5, 200 mM NaCl, 5 mM MgCl2, 1 mM DTT, 0.02% Triton, and 150 ng/µl 3xFlag peptide) by shaking for 30 to 40 min at 4°C.

**Protein stability assay**

Thermostability for purified 3xFlag-LRRK2 was measured by nano-differential scanning fluorimetry using a NanoTemper™ Tycho NT.6 (NanoTemper Technologies, Munich, Germany). Samples were loaded into Tycho NT.6 Capillaries (NanoTemper Technologies, Cat# TY-C001) and the thermal unfolding profiles of WT LRRK2 and the LRKK2 mutants were recorded. Temperature inflection values (Ti) were obtained by automated data analysis. Protein unfolding was followed by the intensity of tryptophan fluorescence at 350 nm/330 nm. Temperature inflection (Ti) is the transition point along the curve that represents to the unfolding transition of the sample. Each variant was recorded in triplicate using three different purification batches.

##### Statistical analyses

For WB quantification, LRRK2 and RAB10 protein levels were normalized against those of GAPDH and the wildtype condition. The phosphorylation levels of LRRK2 at serines 910, 935, and1292 and RAB10 at threonine 73 were normalized against the expression of GAPDH and then evaluated relative to LRRK2 or RAB10 total protein levels, respectively, to obtain the phosphorylation rate. All statistical analysis and graphs were performed using Prism software version 8.0.2 (GraphPad, San Diego, CA, USA). The normality of the data distribution was assessed using the Shapiro-Wilk test. One-way ANOVA and *post hoc* Dunnett's multiple comparison tests were used to determine the difference in the phosphorylation rate between each LRRK2 mutant and the wildtype LRRK2. Paired t-test analyses were performed to compare the phosphorylation rate in the presence of MLi-2 relative to the solvent. Statistical tests were performed at the two-tailed $\alpha$ level of 0.05 for the one-way ANOVA and *post hoc* Dunnett's multiple comparison tests. Statistical tests were performed at the one-tailed $\alpha$ level of 0.05 for paired t-test analysis. All data are expressed as the mean ± standard deviation. P-values are designated as *$p < 0.05$, **$p < 0.01$, ***$p < 0.001$, and ****$p < 0.0001$.

#### DATA AVAILABILITY STATEMENT

The data that support the findings of this study are available from the corresponding authors upon reasonable request.

**Supplementary Table S1. List of primer names and sequences used for site-directed-mutagenesis.**

| Primer name | Sequence |
|---|---|
| LRRK2-H230R-F | GGAATCGCTAGGGAACGTAAACAATGCAGCACATGA |
| LRRK2-H230R-R | TCATGTGCTGCATTGTTTACGTTCCCTAGCGATTCC |
| LRRK2-A1440P-F | ACAGGGGAAGAAGAAGCGCGAGGCTTTATATTGAAGAGCC |
| LRRK2-A1440P-R | GGCTCTTCAATATAAAGCCTGGCGCTTCTTCTTCCCCTGT |
| LRRK2-G2019S-F | GCAGTACTGAGCAATGCTGTAGTCAGCAATCTTTG |
| LRRK2-G2019S-R | CAAAGATTGCTGACTACAGCATTGCTCAGTACTGC |
| LRRK2-R1441G-F | GAAGAAGAAGCGCCAGCCTTTATATTGAAGAGCCAAG |
| LRRK2- R1441G-R | CTTGGCTCTTCAATATAAAGGCTGGCGCTTCTTCTTC |

**Supplementary table S2. Antibodies used in immunohistochemistry experiments**

H+L = Heavy chain + Light chain.

| Antibody | Species | Dilution | Secondary | Source |
|---|---|---|---|---|
| Flag M2 | Mouse | 1/1 000 | Horse anti-mouse IgG Antibody (H+L), Peroxidase | Sigma<br>(Sigma-Aldrich, Saint-Louis, MI, USA) |
| pSer935 LRRK2 | Rabbit | 1/1 000 | Goat anti-rabbit IgG Antibody (H+L), Peroxidase | Abcam<br>(Abcam, Cambridge, UK) |
| pSer910 LRRK2 | Rabbit | 1/1 000 | Goat anti-rabbit IgG Antibody (H+L), Peroxidase | Abcam<br>(Abcam, Cambridge, UK) |
| pSer1292 LRRK2 | Rabbit | 1/500 | Goat anti-rabbit IgG Antibody (H+L), Peroxidase | Abcam<br>(Abcam, Cambridge, UK) |
| RAB10 | Rabbit | 1/1 000 | Goat anti-rabbit IgG Antibody (H+L), Peroxidase | Cell signaling<br>(Cell Signaling, Danvers, MA, USA) |
| pThr73 RAB10 | Rabbit | 1/500 | Goat anti-rabbit IgG Antibody (H+L), Peroxidase | Abcam<br>(Abcam, Cambridge, UK) |
| GAPDH | Rabbit | 1/50 000 | Goat anti-rabbit IgG Antibody (H+L), Peroxidase | Sigma<br>(Sigma-Aldrich, Saint-Louis, |

| | | | | |
|---|---|---|---|---|
| | | | | MI, USA) |
| anti-mouse IgG antibody (H+L) | Horse | 1/50 000 | - | Vector Laboratories<br>(Vector Laboratories, Burlingame, CA, USA) |
| anti-rabbit IgG Antibody (H+L), Peroxidase | Goat | 1/50 000 | - | Vector Laboratories<br>(Vector Laboratories, Burlingame, CA, USA) |

#### SUPPLEMENTARY DATA

##### Clinical phenotypes

###### *Family 1*

A 24-year-old Caucasian woman (IV-3) was referred to the expert center for Parkinson's disease (Lille University Hospital) for a suspicion of PD. For two years, she had had a resting tremor (frequency of 6 to 7 Hz on accelerometric recording) of the left upper limb that was responsive to an acute apomorphine challenge test. Brain MRI and copper and ceruloplasmin levels were normal. She had restless legs syndrome and a medical history of hypothyroidism. There was no professional or environmental toxic exposure but active smoking since the age of 12 years (6 pack-years at first visit) and the drinking of approximately five cups of coffee a day. During examination, she showed typical parkinsonian syndrome, with left-side dominant akinesia, rigidity, and resting tremor, without axial impairment or any red flags. She developed mild impulse control disorders (gambling addiction) with a dopaminergic agonist (rotigotine) and levodopa was introduced at age 35. One year later, she developed motor fluctuations and dyskinesia. She showed left dystonia, anxiety, and abdominal pain during the “off” state, with no axial troubles or cognitive impairment (MOCA = 27/30). At age 36, in the “on” state, her score on the unified Parkinson’s disease rating scale (UPDRS) parts I, II, and III and Hoehn and Yahr score were 2, 5, 5, and 2, respectively. She was treated with levodopa, rasagiline, and rotigotine, with an excellent response for motor signs. She reported a familial history of PD over four generations, with seven affected relatives (Fig. 1B).

Her second cousin (III-1) developed an akineto-rigid syndrome at age 53. The first examination at age 54 showed asymmetrical akinesia, rigidity, and resting tremor, without axial impairment or any red flags (8). He had a history of benign prostatic hypertrophy. There was no professional or environmental exposure, and he did not smoke. At age 64, he underwent bilateral deep brain stimulation of the subthalamic nuclei to control motor fluctuations and dyskinesia. At the last examination (age 66), he presented dysarthria, postural instability, and falls and was apathetic. He was treated with levodopa, rasagiline, tianeptine, and rivastigmine. The MMSE score was 28/30 and the Hoehn and Yahr stage was 3.

***Family 2***

A 67-year-old Caucasian woman (III-29) was referred to the expert center for Parkinson's disease (Lille University Hospital) for a suspicion of PD, having had micrographia and bradykinesia during the previous year. Prodromal features included olfactory disturbances and constipation during the previous several years. She had restless legs syndrome, a history of ankylosing spondylitis, sleep apnea, and high blood pressure. There was no professional or environmental toxic exposure and there was a past history of smoking (17 pack-years). During examination, she showed typical parkinsonian syndrome, with left-side dominant akinesia and rigidity, without resting tremor, axial impairment, or any red flags. The UPDRS part III was 12/108. $^{123}$I-FP-CIT single-photon emission computed tomography (DatSCAN imaging) indicated decreased striatal uptake, predominating on the right-side. She was treated with levodopa and a dopaminergic agonist (rotigotine) with a good response for motor signs. Three years later, she developed urinary urgency, impulse control disorders (binge eating, hobbyism), wearing-off, motor fluctuations, and mild dyskinesia. She showed no dystonia, axial impairment, or cognitive disturbances. She reported a familial history of PD over three generations, with 11 affected relatives (Fig. 1C).

Patient III.1, a Caucasian man, developed slowing of gait around the age of 73 years. Anxiety and insomnia had been present for several years before the first motor symptoms, but there were no other prodromal non-motor features. He had no history of smoking or toxin exposure. Clinical exam at the age of 76 showed general bradykinesia, mild rigidity in the four limbs and mild rest tremor in both feet. He walked with small steps and arm swing was reduced bilaterally. Brain MRI showed no significant ischemic or other lesions. $^{123}$I-FP-CIT SPECT showed reduced striatal uptake, especially in the right putamen. He showed a good response to levodopa. He did not develop motor fluctuations or dyskinesias. Insomnia and anxiety remained prominent. There were no psychotic or significant cognitive symptoms. At the age of 81, Mini Mental State Examination (MMSE) score was 28/30. At the age of 82, UPDRS III score ("on" levodopa) was 26.

Patient III.2 was a Caucasian woman who first noticed dragging of her right foot around the age of 62 years. She had suffered from depressive symptoms for several years, but there were no other prodromal non-motor features. There was no history of smoking or toxic exposure. Clinical exam at the age of 64 showed right-sided bradykinesia and rigidity and rest tremor of the right hand. Brain MRI showed no significant abnormalities. $^{123}$I-FP-CIT SPECT showed reduced striatal uptake, especially in the left putamen. Levodopa and ropinirole had a favorable effect on her motor symptoms. She gradually developed motor fluctuations and levodopa-induced dyskinesias. At the age of 74, treatment with levodopa-carbidopa intestinal gel (LCIG) was initiated, resulting in increased "on" time. She continued to have depressive symptoms and developed chronic constipation. Gait problems worsened significantly around the age of 80 years, leading to repeated falls and use of a rollator. There were no psychotic symptoms or dementia. At the age of 81, MMSE score was 26/30, scores on UPDRS part I, II, III (“on” LCIG) and IV were 1, 22, 30 and 4, respectively, and Hoehn-Yahr score (“on” LCIG) was 4.

**Thermal stability**

Thermodynamic stability is a fundamental property of proteins that significantly influences protein structure, function, expression, and solubility. Here, we assessed how the identified point mutations affect the stability and folding of the LRRK2 protein.

The H230R and A1440P variant proteins did not show notable differences in their respective profiles or Ti relative to the WT protein (Supplementary Figure S3A-B). The Ti was higher for G2019S (64.6°C ± 1.45; $p = 0.047$) and R1441G (66.5°C ± 0.71; $p < 0.0001$) than for the WT protein (63.1°C ± 0.14). The initial ratio, which indicates the level of folding at basal condition, was similar for H230R (0.34 ± 0.019; $p = 0.97$) and lower for A1440P (0.31 ± 0.011; $p = 0.036$), G2019S (0.3 ± 0.019; $p = 0.033$) and R1441G (0.3 ± 0.015; $p < 0.02$) than for the WT protein (0.34 ± 0.004) (Supplementary Figure S3C). There was no difference in the Δ Ratio (Initial ratio – final ratio). Our results suggest that these proteins are correctly folded and stable, despite the amino-acid substitutions.

**Supplementary Figure S1. Reduced phosphorylation of LRRK2 after MLi-2 treatment.**

Representative western blot and quantification of LRRK2 phosphorylation at serines 910 **(A-B)**, 935 **(C-D)**, and 1292 **(E-F)** after DMSO or DMSO + Mli-2 treatment in WT and LRRK2 mutants. Error bars indicate the standard deviation of replicates (n = 3). kDa = Kilodalton. *$p < 0.05$, **$p < 0.01$, ***$p < 0.001$.

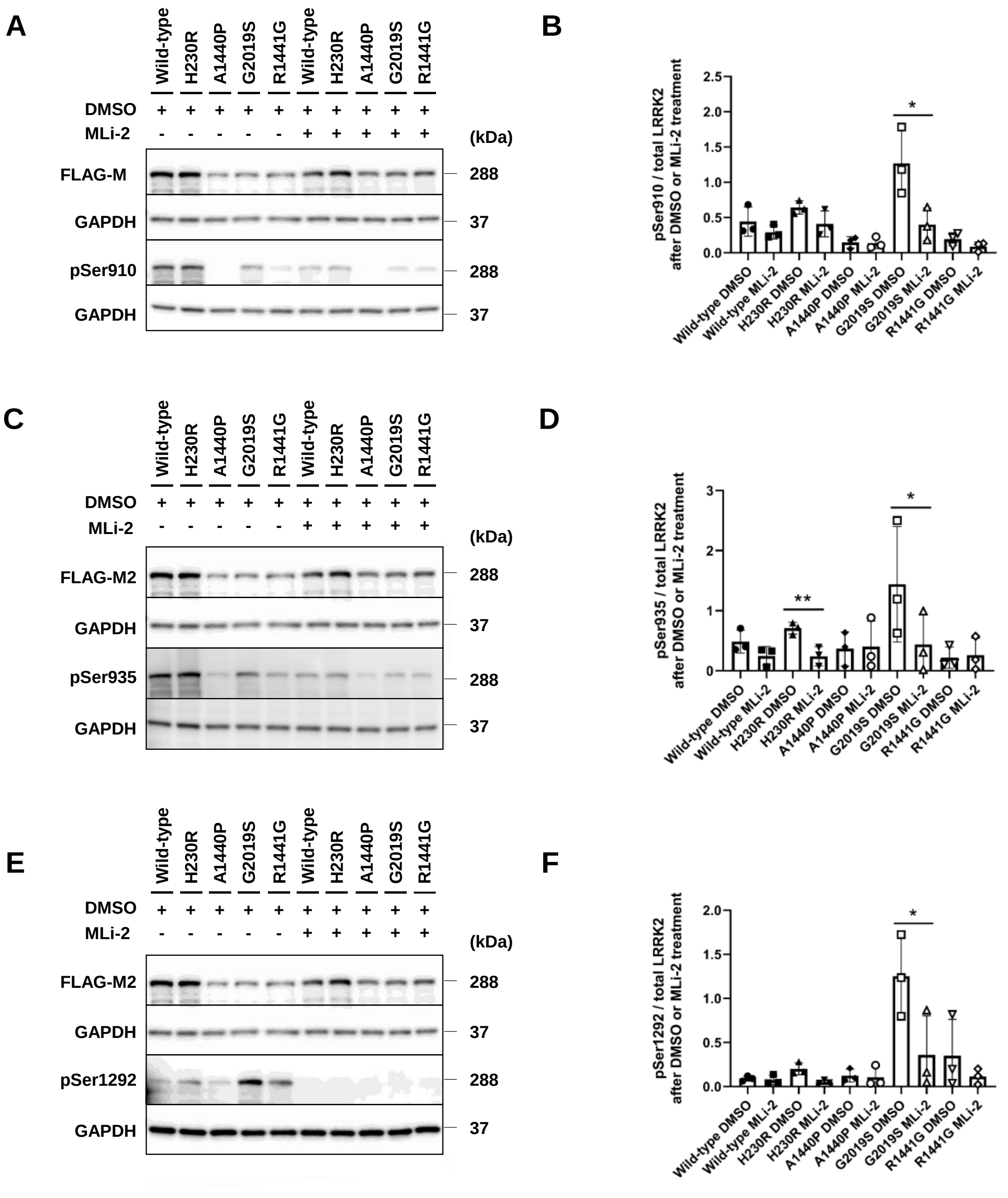

**Supplementary Figure S2. Inhibition of LRRK2 kinase activity**

Representative western blot **(A)** and quantification **(B)** of RAB10 phosphorylation at threonine 73 after MLi-2 treatment (100 nM, 1h) or DMSO treatment (0.01 %, 1h), as a negative control. Error bars indicate the standard deviation of replicates (n = 3). No statistical test was performed. kDa = Kilodalton.

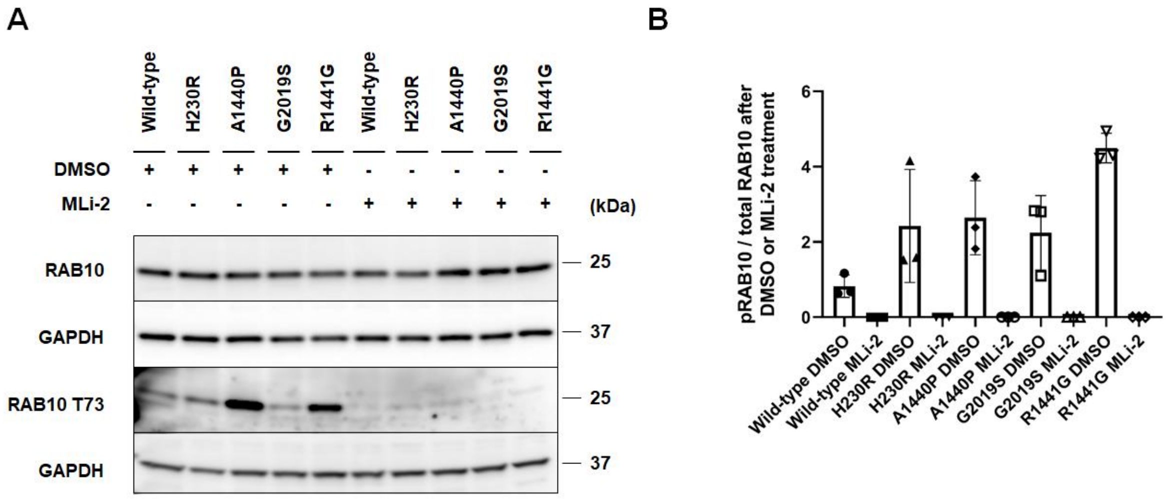

**Supplementary Figure S3. Thermodynamic stability of LRRK2 variants.**

**A.** Thermal degradation profile of LRRK2 proteins. Vertical lines indicate the Ti. **B.** Quantification of LRRK2 protein temperature inflection values. Error bars indicate the standard deviation of replicates (n = 3). **C.** Quantification of the initial ratio of fluorescence intensity for the LRRK2 protein at 350 nm/330 nm. *$p < 0.05$, **$p < 0.01$, ***$p < 0.001$, ****$p < 0.001$.

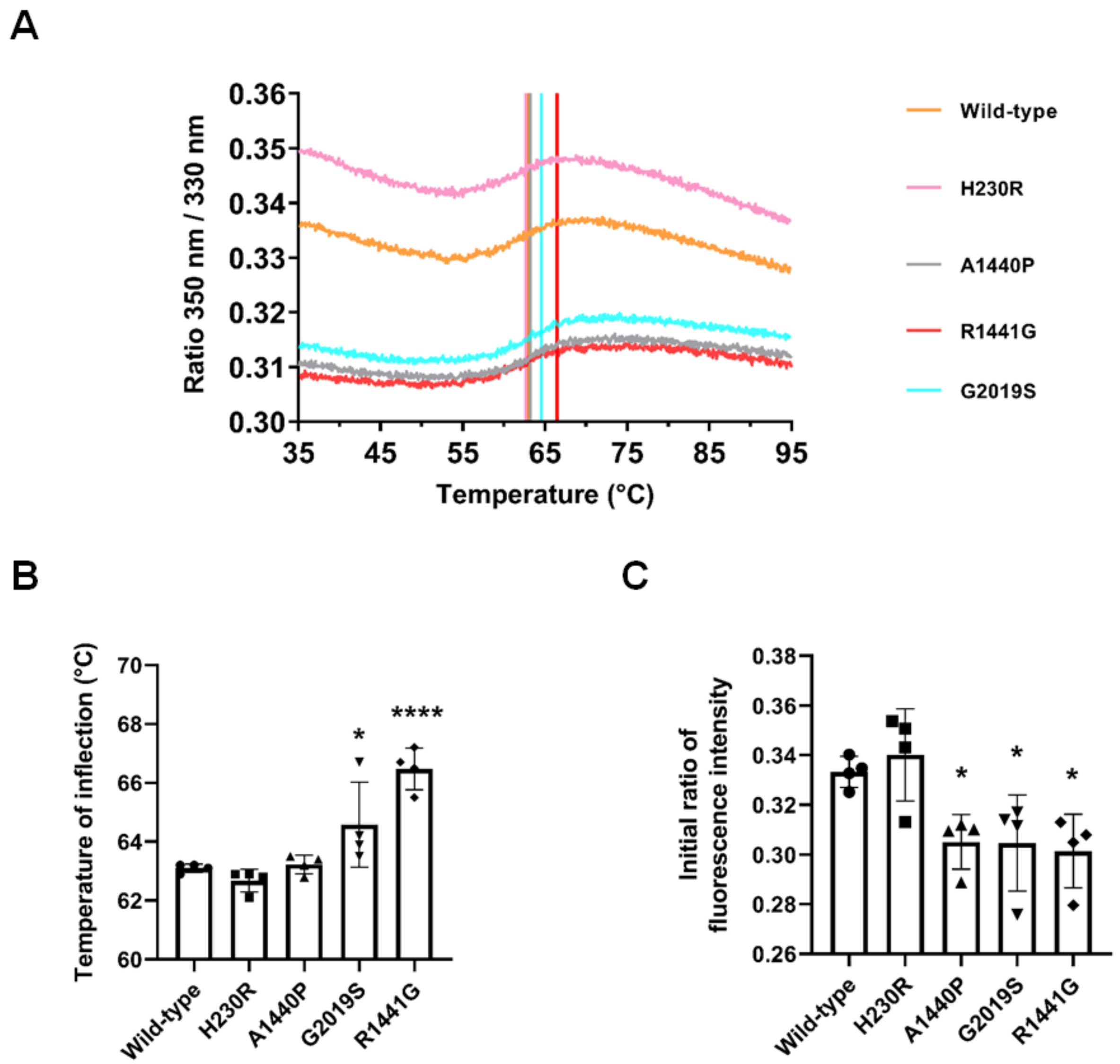
